# Sofosbuvir protects human brain organoids against SARS-CoV-2

**DOI:** 10.1101/2020.05.30.125856

**Authors:** Pinar Mesci, Angela Macia, Aurian Saleh, Laura Martin-Sancho, Xin Yin, Cedric Snethlage, Simoni Avansini, Sumit K. Chanda, Alysson Muotri

## Abstract

COVID-19 was rapidly declared a pandemic by the World Health Organization, only three months after the initial outbreak in Wuhan, China. Early clinical care mainly focused on respiratory illnesses. However, a variety of neurological manifestations in both adults and newborns are also emerging. To determine whether SARS-CoV-2 could target the human brain, we infected iPSC-derived human brain organoids. Our findings show that SARS-CoV-2 was able to infect and kill neural cells, including cortical neurons. This phenotype was accompanied by impaired synaptogenesis. Finally, Sofosbuvir, an FDA-approved antiviral drug, was able to rescue these alterations. Given that there are currently no vaccine or antiviral treatments available, urgent therapies are needed. Our findings put Sofosbuvir forward as a potential treatment to alleviate COVID-19-related neurological symptoms.

**One Sentence Summary:** SARS-CoV-2 infection causes neuronal death and impaired synaptogenesis, both rescued by Sofosbuvir treatment.

## Main Text

Coronavirus disease 2019 (COVID-19) caused by the Severe Acute Respiratory Syndrome Coronavirus 2 or SARS-CoV-2, was declared a pandemic by the World Health Organization on March 11, 2020. The outbreak first began in Wuhan, China, in December 2019 and quickly spread to 216 countries in 5 months with more than five million cases and more than 330,000 confirmed deaths worldwide. There are now more than one and a half million confirmed cases in the United States, with deaths nearing 100,000 as of May 24, 2020. With no vaccines, approved treatments, and a high infectivity rate, COVID-19 quickly became a public health emergency worldwide, leading to lockdowns in many areas of the world severely affecting daily life.

As the situation has been developing rapidly, early studies mainly focused on the respiratory component of COVID-19 disease. However, as more cases appear, other COVID-19-related clinical manifestations have been reported. Indeed, some COVID-19 adult patients presented with several other neurological symptoms including stroke, hallucinations, epilepsy, encephalopathy, anosmia or ageusia, suggesting that SARS-CoV-2 could impact the central nervous system (*1*–*6*). Other neurological manifestations will likely surface in the near future. Miscarriages and premature births to COVID-19-positive pregnant women have also been reported (*7*–*9*). Recently, a study showed that placentas from COVID-19-positive pregnant women displayed injury (*10*). Finally, SARS-CoV-2 was also detected in semen(*11*). Altogether, these findings support the possibility of vertical transmission of the virus to the fetus and potentially affect brain development(*12*). Supporting this hypothesis, babies born to SARS-CoV-2-positive mothers have shown several inflammatory symptoms such as neonatal sepsis, rashes, eye infections with long-term impacts remaining unclear (*8*). However, there is no experimental evidence that demonstrates that the Sars-CoV-2 virus can infect and impact human brain cells.

Our laboratory previously revealed a causative link between the circulating Brazilian Zika virus and the severe microcephaly observed in babies born from infected mothers using human induced pluripotent stem cell (hiPSC)-derived brain organoids (*13*). Human brain organoids are scaled down, three-dimensional models of the brain, recapitulating several molecular and cellular aspects of human embryonic and fetal stages (*14*). Brain cortical functional organoids closely mimicked the early stages of human neurodevelopment and network formation (*15*). Here, we tested whether SARS-CoV-2 could infect the developing human brain and whether the FDA-approved antiviral drug, Sofosbuvir, could be considered a potential treatment.

Angiotensin-converting enzyme-2 (ACE2) is a critical receptor for SARS-CoV-2; hence its expression has been used to predict the potential vulnerability of different cell types (*16*). We generated eight-week-old human brain cortical organoids from healthy donors that contain neural precursor cells (Nestin^+^, NPC), cortical neurons (MAP2^+^), and astrocytes (GFAP^+^) (Fig. 1A). We found that ACE2 receptors are expressed in human iPSC-derived brain cortical organoids, neurons, astrocytes but not microglia (Fig. 1B). Next, brain cortical organoids were infected with a SARS-CoV-2 isolate obtained from a patient in Washington State at a multiplicity of infection (MOI) of 2.5. Brain cortical organoids were then assessed by immunolabeling one-week postinfection. Both TUNEL and cleaved caspase 3 (CC3)-integrated signal density were significantly increased by 20- and 5-fold respectively in organoids infected with SARS-CoV-2 compared to mock-infected ones (Fig.1C-D). TUNEL and CC3 increased signals were accompanied by a rise in the staining of the SARS-CoV-2 virus by 10-fold compared to mock conditions (Fig.1E).

We next examined which cell type was preferentially targeted and susceptible to infection with SARS-CoV-2 (Fig.1–2). Eight-week-old organoids contain mostly NPC, neurons, and few astrocytes (*15*), (Fig 1A). Therefore, we studied cell death within each of these cell populations. NPCs showed a 30-fold increase in cell death measured by TUNEL (Fig.1F-G), accompanied by an accumulation of SARS-CoV-2 within Nestin^+^ NPCs (Fig. 1H). Next, we studied cell death and viral accumulation in GFAP+ astrocytes (Fig.2A-C). Upon infection with SARS-CoV-2, astrocytes had a 4-fold increase in cell death but no viral accumulation was observed (Fig.2A-C). Finally, we assessed whether MAP2^+^ cortical neurons could be the targets of SARS-CoV-2. Upon infection with SARS-CoV-2, we observed a 7-fold increase in cell death in MAP2^+^ cells, accompanied by an increase in the number of MAP2 and SARS-CoV-2 double-positive cells (Fig.2D-F).

**Figure 1.**
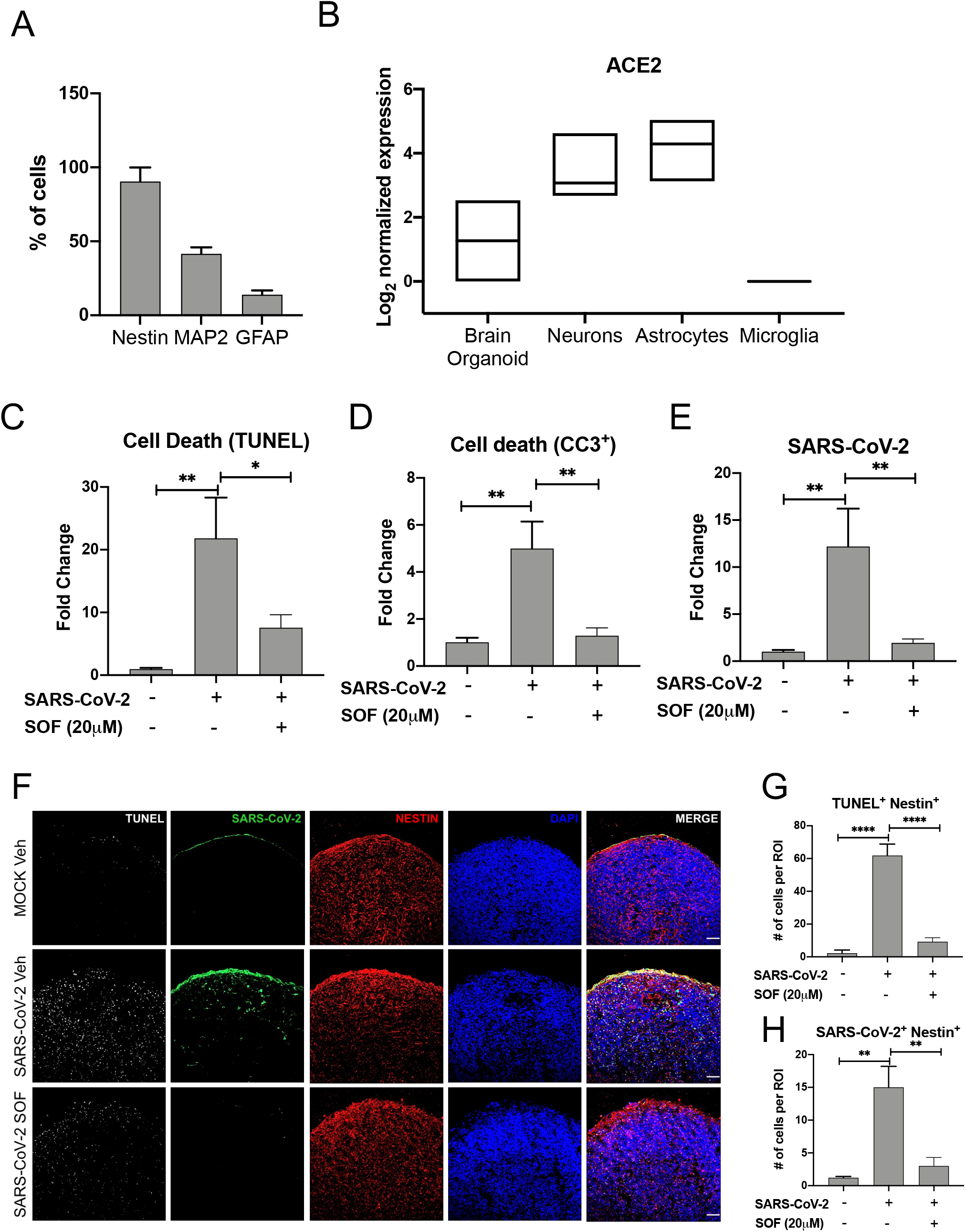
Impact of SARS-Cov-2 infection in cortical organoids. **A**. Percentage of Nestin, MAP2, and GFAP expressing cells in 3D cortical brain organoids. **B.** Normalized mRNA expression of *ACE2* in iPSC-derived brain organoids, neurons, astrocytes, and microglia. **C-D.** Quantification of TUNEL and CC3 integrated densities. Treatment with 20μM Sofosbuvir (SOF) significantly reduces TUNEL and CC3 expression in SARS-CoV-2 infected organoids, n=5-6 biological replicates per condition. Data is normalized to mock vehicle conditions. Error bars represent S.E.M. *p<0.05, **p<0.01, One-way ANOVA and Tukey’s multiple comparisons test. **E.** Day-52 cortical organoids support SARS-CoV-2 infection at MOI 2.5. Measurement of SARS-CoV-2 integrated signal density upon vehicle (Veh) or treatment with 20μM Sofosbuvir (SOF). **F.** Immunolabelling of TUNEL (white), SARS-CoV-2 (green) and Nestin (red) in mock, infected, and SOF treated organoid sections. Scale bar, 50 μm. **G.** Quantification of TUNEL^+^ within Nestin^+^ neural precursor cells (NPC) following infection with SARS-CoV-2 per region of interest (ROI), n=5-6 biological replicates per condition. Data is normalized to mock vehicle conditions. Error bars represent S.E.M. ****p< 0.0001, One-way ANOVA and Tukey’s multiple comparisons test. **H.** Quantification of SARS-CoV-2^+^ within Nestin^+^ cells following infection with SARS-CoV-2 per ROI, n=5-6 biological replicates per condition. Data is normalized to mock vehicle conditions. Error bars represent SEM,**p<0.01, One-way ANOVA and Tukey’s multiple comparisons tests.

**Figure 2:**
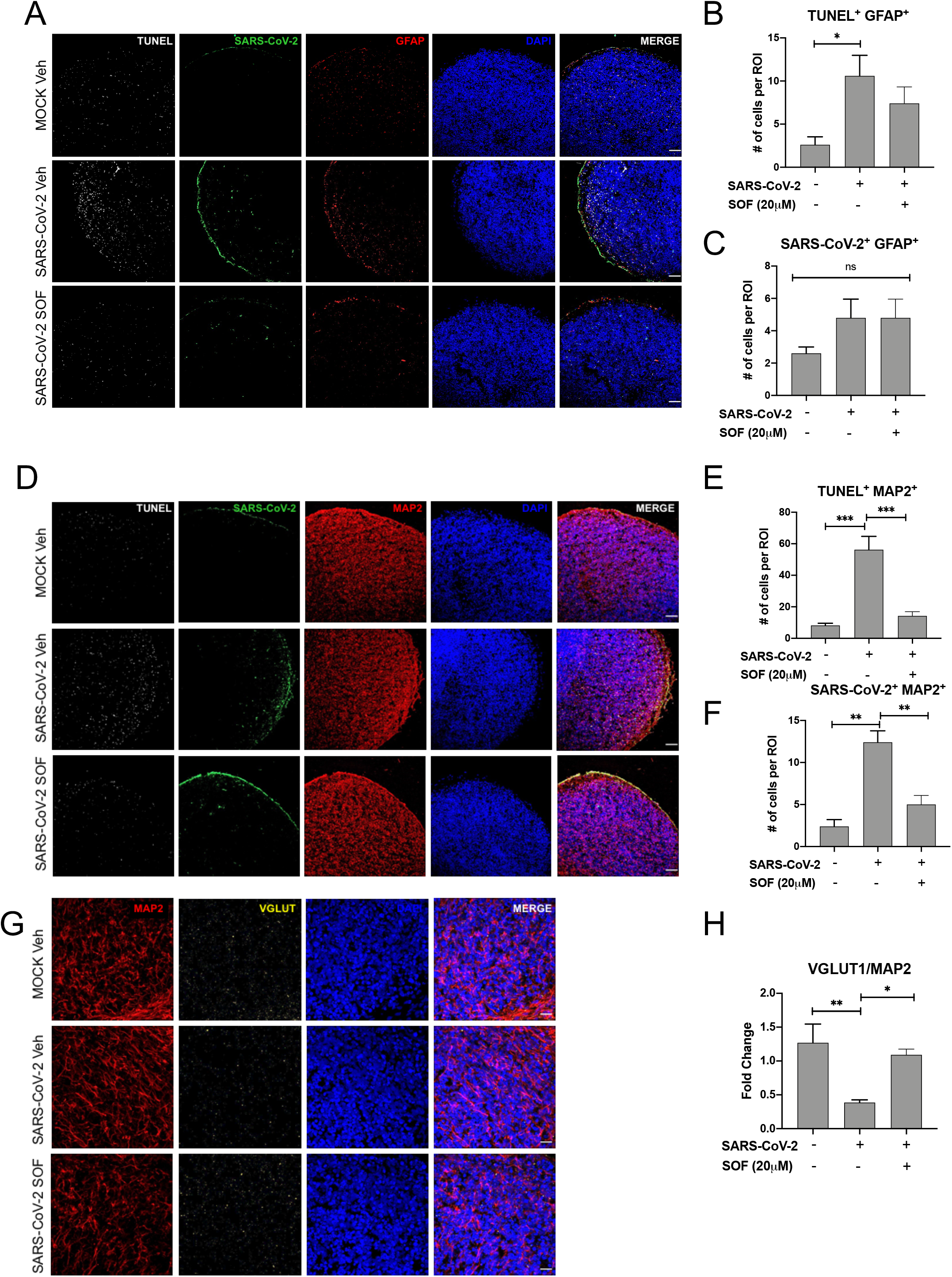
SARS-CoV-2 induces cortical neuronal death and decrease excitatory synapses. **A-D-G.** Immunolabeling of mock, infected, and infected, and SOF treated organoid sections stained for TUNEL (white), SARS-CoV-2 (green), GFAP (red), MAP2 (red), and vGLUT1 (in yellow) respectively confocal microscopy. Scale bar, 50 μm for A-D and 20 μm for G, n=5-6 biological replicates per condition. **B-C** Quantification of the frequency of GFAP expressing cells (red) colocalizing with TUNEL (white) and SARS-CoV-2 positive cells (green) per ROI n=5-6 biological replicates per condition. Data is normalized to mock vehicle conditions. Bars represent mean, error bars represent SEM *p<0.05. One-way ANOVA and Tukey’s multiple comparisons test. **E-F:** Quantification of the frequency of MAP2 expressing cells (red) colocalizing with TUNEL (white) and SARS-CoV-2 positive cells (green) per ROI, n=5-6 biological replicates per condition. Data is normalized to mock vehicle conditions. Bars represent mean, error bars represent SEM **p<0.01, ***p<0.001, One-way ANOVA and Tukey’s multiple comparisons test. **H.** Quantification of vGLUT1 positive cells (yellow) within MAP2^+^ cortical neurons with or without infection with SARS-CoV-2 per ROI, MOI 2.5, n=5 biological replicates per condition. Data is normalized to mock vehicle conditions. Bars represent mean. Error bars represent S.E.M. *p<0.05, **p<0.01. One-way ANOVA and Tukey’s multiple comparisons test.

Given that SARS-CoV-2 could infect MAP2^+^ post-mitotic neurons, we wondered whether the infection could also affect synaptogenesis. To assess the impact on synaptogenesis, we quantified the number of excitatory synapses in brain organoids using vGLUT1 antibodies (Fig2G-H). The pre-synaptic protein vGLUT1 was significantly decreased by 70 % upon infection with SARS-CoV-2 compared to mock-conditions (Fig.2H).

We previously used brain organoids to repurpose drugs for several neurological conditions (*17*–*20*). Here, we tested Sofosbuvir efficacy in blocking SARS-CoV-2-induced neurotoxicity. Sofosbuvir (Sovaldi, Gilead Sciences) is an FDA-approved anti-hepatitis C treatment that blocks HCV replication by inhibiting its RNA-dependent RNA polymerase (RdRp). Sofosbuvir can also suppress other viral families of single-stranded, positive-sense RNA viruses, of which Coronaviruses belong to (*19*, *21*). The SARS-CoV-2 RdRp shares high sequence and structural homology with HCV, supporting the likelihood of inhibiting SARS-CoV-2 with similar efficiency, therefore, we hypothesized that Sofosbuvir could inhibit SARS-CoV-2 RdRp, preventing neuronal death. We determined the safe treatment dosage for brain organoids for Sofosbuvir based on our previous study (*19*). We found that treating the organoids twice a week with 20μM Sofosbuvir post-infection, successfully decreased the amount of neuronal cell death and viral accumulation in brain organoids (Fig1C-H, Fig2). Moreover, we also show that Sofosbuvir restored vGLUT1 protein expression in synaptic puncta (Fig2G, H). The immediate and long-term lasting neurological and neuropsychiatric sequelae of COVID-19 is currently surfacing. It might take several years to document the survivor’s cognitive and mental health burden of recovered COVID-19 cases (*22*). Our findings show that SARS-CoV-2 can infect different cell types in human cortical organoids, including postmitotic neurons. Moreover, SARS-CoV-2 rapidly decreased the amount of the excitatory synapses in cortical neurons, revealing a potential mechanism for the associated neurological symptoms with COVID-19. Our data provide direct experimental evidence that the developing human brain is susceptible to SARS-CoV-2 infections, likely leading to long-term impairments in neuronal function in babies born from COVID-19-positive mothers. In the absence of a vaccine, efficient drug treatment for COVID-19 is urgently in need. We show that the observed neurological impairments in infected brain organoids are rescued by Sofosbuvir treatment, a brain-penetrant FDA-approved drug (*23*). Supporting our findings, Sofosbuvir has been pointed as a potential treatment against COVID-19 based on *in silico* modeling (*24*). Thus, treatment with Sofosbuvir could arrest or prevent the development of neurological symptoms in COVID-19 adults. Sofosbuvir is also an option to block vertical transmission from SARS-CoV-2-infected pregnantwomen for whom prevention is no longer an option. Although further clinical studies are needed, we provide initial evidence that Sofosbuvir is an immediate candidate to pharmacologically treat neurological manifestations related to COVID-19.

## Author Contribution

P.M., A.M., A.S., and A.R.M. conceived this study. P.M., A.M., A.S. designed and performed all the experiments. X.Y. L.M.S. performed the infections with SARS-CoV-2. C.S. and S.A. generated brain cortical organoids. C.S. helped with sectioning and immunolabelling. P.M., A.M., A.S. interpreted the results, and P.M. wrote the manuscript. All authors revised, contributed, and approved the final manuscript.

## Data and materials availability

All data is available in the main text or the supplementary materials.

## Materials and Methods

### Viral Isolate

SARS-CoV-2 USA-WA1/2020 strain, isolated from an oropharyngeal swab from a patient with a respiratory illness who developed clinical disease (COVID-19) in January 2020 in Washington, USA, was obtained from BEI Resources (NR-52281). The virus was propagated in Vero E6 cells (ATCC^®^ CRL-1586TM) transfected with exogenous human ACE2 and TMPRSS2. Virus titers were determined by plaque assay performed on Vero E6 cells. All experiments involving live SARS-CoV-2 followed the approved standard operating procedures of the Biosafety Level 3 facility at the Sanford Burnham Prebys Medical Discovery Institute.

### RNAseq analyses

Brain cortical organoids, hiPSC-derived microglia, astrocyte and neurons were generated as previously described (*15*, *17*, *25*) and total RNA was extracted using RNeasy Qiagen Mini kit according to manufacturer’s instructions. RNAseq was performed in a Illumina HiSeq4000. Data was analyzed by Rosalind ((https://rosalind.onramp.bio/), an online platform for data analyses, as previously described (*18*).

### Cell lines

Vero E6 cells were maintained in Dulbecco’s modified eagle medium (DMEM, Gibco) supplemented with 10 % heat-inactivated fetal bovine serum (FBS, Gibco), 50 U/mL penicillin, 50 μg/mL streptomycin, 1 mM sodium pyruvate (Gibco), 10 mM HEPES (Gibco), and 1X MEM non-essential amino acids solution (Gibco).

Donated healthy fibroblasts were obtained via skin biopsies from patients after informed consent was appropriately given under protocols approved by the University of California, San Diego Institutional Review Board (#141223ZF). All experiments were approved and performed under the Institutional Review Boards (IRB) and Embryonic Stem Cell Research Oversight (ESCRO) guidelines and regulations.

### Cell culture

iPSCs were cultured and manually passaged onto Matrigel-coated (Corning) and fed daily with mTESR1 (StemCell Technologies).

iPSCs-derived brain cortical organoids were differentiated as previously described (Trujillo et al., 2019). Briefly, iPSCs were dissociated using a 1:1 Dulbecco’s phosphate-buffered saline (DPBS, ThermoFisher) and StemPro Accutase (ThermoFisher) solution. Cells were centrifuged and resuspended in mTeSR1 supplemented with 10 μM SB431542 (SB; Stemgent) and 1 μM Dorsomorphin (R&D Systems). 4×106 cells were transferred to one well of a 6-well plate and kept in suspension under rotation (95 rpm) for 24 hours with 5 μM of ROCK inhibitor (Y-27632; Calbiochem). Forty-eight hours later, media was substituted by the neural induction media consisting of DMEM/F12 (Life Technologies), 1% Glutamax (Life Technologies), 1% N2 Neuroplex (Gemini Bio), 1% non-essential amino acids (NEAA, Gibco), 1% Pen-Strep (ThermoFisher), 10μM SB431542 and 1μM of Dorsomorphin. The media was changed every other day for seven days. Media was substituted for neural proliferation media consisting of Neurobasal media (Life Technologies), 2% Gem21 Neuroplex, 1% non-essential amino acids, 1% Glutamax. And 20 ng/mL basic fibroblast growth factor (bFGF; Life Technologies). The media was changed daily for 7 days, followed by another seven days of neural proliferation media supplemented with 20 ng/mL of epidermal growth factor (EGF, Peprotech). Neuronal Maturation step was achieved by changing the media to Neurobasal with GlutaMAX, 1% Gem21 NeuroPlex (Gemini Bio), 1% NEAA and 1% PS; supplemented with 10 ng/mL of BDNF, 10 ng/mL of GDNF, 10 ng/mL of NT-3 (PeproTech), 200 mM L-ascorbic acid and 1 mM dibutyryl-cAMP (Sigma-Aldrich). The media was replaced every other day for seven more days. We executed cortical organoid experiments using maintenance media: Neurobasal with GlutaMAX, 1% Gem21 NeuroPlex (Gemini Bio), 1% NEAA, and 1% PS.

All the cell lines tested negative for mycoplasma contamination. All cell lines used have been authenticated.

### *In vitro* infection

Brain cortical organoids were infected with 750,000 PFU that correspond to a MOI of 2.5 considering an average of 300,000 cells per organoid on day 52 of differentiation and left in culture for three days in maintenance media. Fresh media was added atop and kept in suspension for another four days. 20μM of Sofosbuvir (SOF; Acme Bioscience AB3793) added to cell culture supernatant at the desired concentration on the day of infection and 4 days post-infection. Brain cortical organoids were fixed 7 days post-infection for 48 hours with a 4% paraformaldehyde (PFA) solution, and further embedded in 30% sucrose for 24 hours. Tissue was placed in a mold for cryosectioning and covered with a layer of optimal cutting temperature (CRYO-OCT; VWR) compound to prevent freeze-drying and store the rest of the sample at −80°C. Samples were sliced in 20μm cryosections using (Leica CM3050) and prepared for immunostaining (see below for further details).

### Immunofluorescence and imaging analyses

Brain cortical organoids were fixed with 4% paraformaldehyde for 48 hours at 4°C. Next, samples were permeabilized in 1xPBS (Corning) containing 0.1% (v/v) Triton X-100 for 10 minutes. Fixed cells were next incubated with blocking solution for 1 hour [3% Bovine Serum Albumin (BSA); (Gemini) in 1xPBS]. Primary antibodies were diluted with blocking solution and incubated with cells overnight at 4°C: Cleaved caspase-3 (rabbit, Cell Signaling #9661, 1:500), Anti-Nestin (mouse, Anti-Nestin antibody [10C2]; 1:1000), Anti-Ki67 antibody (rabbit, Abcam ab15580, 1:300), Anti-MAP2 antibody (Chicken, Abcam ab5392, 1:1000), Anti-VGlUT1 [317D4] (mouse, Synaptic Systems #135311, 1:500), Anti Homer 1 (VesL 1, Syn 47) (rabbit, Synaptic Systems #160003, 1:500), anti-GFAP (chicken, Abcam ab4674, 1:1000). SARS-CoV-2 (2019-nCoV) Nucleoprotein / NP Antibody (rabbit Mab, Sino Biological #40143-R019, 1: 2000) was incubated diluted with blocking solution and incubated with cells for 20 minutes at room temperature. Cells were then washed twice with 1xPBS and incubated with the secondary antibody for 30 minutes at room temperature. Secondary antibodies (all conjugated to Alexa Fluor 488, 555 and 647) were purchased from Life Technologies and used at a 1:1000 dilution. After the 30 minutes incubation, samples were washed twice (1xPBS), incubated for 5 minutes with fluorescent nuclear DAPI stain (VWR; 1:10000), and mounted with Slow fade gold antifade reagent (Life Technologies). Samples were imaged using an Axio Observer Z1 Microscope with ApoTome (Zeiss). For TUNEL analysis, samples were fixed with 4% paraformaldehyde and permeabilized with 0.25% Triton X-100 for 15 minutes and then stained for TUNEL following manufacturer’s instructions (Click-iT TUNEL assay kit from Life Technologies). Cells were blocked with 3% BSA for 1 hour and then incubated in primary and secondaries, as shown above. Images were blindly collected using an Axio Observer Z1 Microscope with ApoTome (Zeiss) and analyzed with ImageJ software. To calculate the integrated density, the channels were split and the fluorescence corresponding to each staining was measured by Image J software.

### Statistical analyses

Results were analyzed using Prism Software (version 6, GraphPad, USA). Statistical significance was determined using one-way ANOVA tests followed by Tukey or Sidak multiple comparisons tests to compare different groups with one variable using a p< 0.05. The reported values are means ± SEM, as mentioned in relevant figure captions. Sample sizes, n, reported in figure legends.

